# RNA polymerase II pausing contributes to maintain chromatin organization in erythrocytes

**DOI:** 10.1101/2022.06.16.496295

**Authors:** Penagos-Puig Andrés, Claudio-Galeana Sherlyn, Stephenson-Gussinye Aura, Jácome-López Karina, Aguilar-Lomas Amaury, Pérez-Molina Rosario, Furlan-Magaril Mayra

**Affiliations:** Departamento de Genética Molecular, Instituto de Fisiología Celular, Universidad Nacional Autónoma de México, Mexico City, Z.C.04510

## Abstract

Chicken erythrocytes are nucleated cells often referred to as transcriptionally inactive, although the epigenetic changes and chromatin remodeling that mediate transcriptional repression and the extent of gene silencing during avian terminal erythroid differentiation are not fully understood. Here we characterized the changes in gene expression, chromatin accessibility, genome organization, and chromatin nuclear disposition during the terminal stages of erythropoiesis in chicken and found a complex chromatin reorganization at different genomic scales. We identified a robust decrease in transcription in erythrocytes. Nevertheless, a set of genes maintains their expression in erythrocytes, including genes involved in RNA pol II promoter-proximal pausing. Erythrocytes exhibit an inverted nuclear architecture and reposition euchromatin towards the nuclear periphery together with the paused RNA polymerase. In erythrocytes, chromatin domains are partially lost genome-wide except at mini domains retained around paused promoters. Our results suggest that promoter-proximal pausing of the RNA pol II participates in the transcriptional regulation of the erythroid genome and highlight the role of RNA polymerase in the maintenance of local chromatin organization.

## Introduction

Red blood cells (RBC) are highly specialized cells that transport oxygen from the lungs to all tissues in the body. In mammals, terminal stages of erythroid differentiation involve a reduction in nuclear size, extensive chromatin condensation mediated by HDACs^1-3^, and remodeling of the nuclear envelope^4^ followed by the expulsion of the nucleus from the erythroblasts^5^. Erythropoiesis in non-mammalian vertebrates like birds however, gives rise to circulating erythrocytes that retain their nucleus in a highly compacted state^6^.

Genome-wide chromatin condensation and gene expression silencing in nucleated RBC in chicken is promoted by the incorporation of the erythroid-specific linker histone H5 into chromatin^7-9^. In fact, chicken RBC undergo a dramatic drop in RNA synthesis and are regarded as transcriptionally inert cells^10-12^, although the exact extent of this repression has not been studied. The chicken nucleated erythrocyte has been useful in the study of chromatin structure and transcriptional regulation^13^. However, the molecular mechanisms that mediate the extensive silencing of the erythroid genome are largely unknown and a study at high resolution of the remodeling of the chromatin and genome organization during this process is still lacking.

The three-dimensional organization of chromosomes in the interphase nucleus serves as a scaffold of chromatin interactions that ensures the execution of complex transcriptional programs^14^. In the last decades, Hi-C experiments unveiled multiple levels of genome organization occurring at different genomic scales^15^. At the megabase scale, the genome is partitioned into two compartments, A and B, which correspond to predominantly active and inactive regions of the genome respectively^16^. At the sub-megabase scale, topologically associated domains (TADs) represent frequently interacting regions that are partly insulated from other neighboring sequences^17,18^. TADs enable proper promoter-enhancer communication inside the domain and prevent interference by regulatory elements outside the TAD and disruption of TAD boundaries leads to aberrant gene activation and disease ^19-21^, highlighting the role of TADs in the spatio-temporal regulation of gene expression. In vertebrates, TADs are formed by the loop-extrusion activity of the cohesin complex and CTCF^22-24^ although the presence of CTCF-independent TADs has been reported^22,25^. The existence of other factors that contribute to TAD formation is still under investigation and it has been suggested that transcription or at least certain components of the transcriptional machinery might mediate TAD establishment^26-28^.

Here, we studied the changes in chromatin accessibility and genome organization that take place in the terminal erythroid differentiation of chicken RBC. We show that chicken erythrocytes undergo a complex multilayered remodeling of chromatin organization. First, we show that chicken RBC have a unique inverted nuclear architecture and relocate their open chromatin towards the nuclear periphery. Despite a genome-wide reduction in gene expression, chicken RBC retain transcription of erythroid-function related genes and genes involved in RNA pol II promoter-proximal pausing. Furthermore, chromatin accessibility is maintained at promoters of inactive genes and colocalizes with the presence of RNA polymerase II in a paused state. Hi-C experiments show a strengthening of chromatin compartmentalization consistent with confocal microscopy imaging of euchromatin, transcription and heterochromatin foci. Moreover, the erythroid genomes are largely devoid of TADs but retain local organization at mini domains around paused genes. Together, our data provide an in-depth characterization of chromatin remodeling during erythropoiesis in chicken. We identified a type of domain organization that persist in absence of transcription and propose that RNA polymerase II binding might be sufficient to maintain local organization.

## Results

### Chromatin accessibility is restricted to the nuclear periphery in aRBC

Chicken erythrocytes are regarded as mostly transcriptionally inactive cells that exhibit a dramatic drop in RNA synthesis ^29^. To investigate the chromatin remodeling process during terminal erythroid differentiation in chicken we obtained primary embryonic (eBRC) and adult (aRBC) red blood cells from 10-day-old chicken embryos and 30 weeks old hens, respectively. We also analyzed HD3 cells^30^ that correspond to an early erythroblast and primary fibroblasts were also used as a non-erythroid cell control.

First, we sought to explore the effect of this genome-wide silencing process on chromatin accessibility and nuclear architecture by labeling and imaging accessible chromatin using ATAC-see^31^ (Fig. 1a; Extended data 1a). ATAC-see signal intensity in eRBC and aRBC is lower than in erythroblasts suggesting an overall decrease in accessibility during erythroid terminal differentiation (Fig. 1a; Extended data 1b). However, chicken erythrocytes retain accessible chromatin even in absence of transcription.

**Fig. 1.**
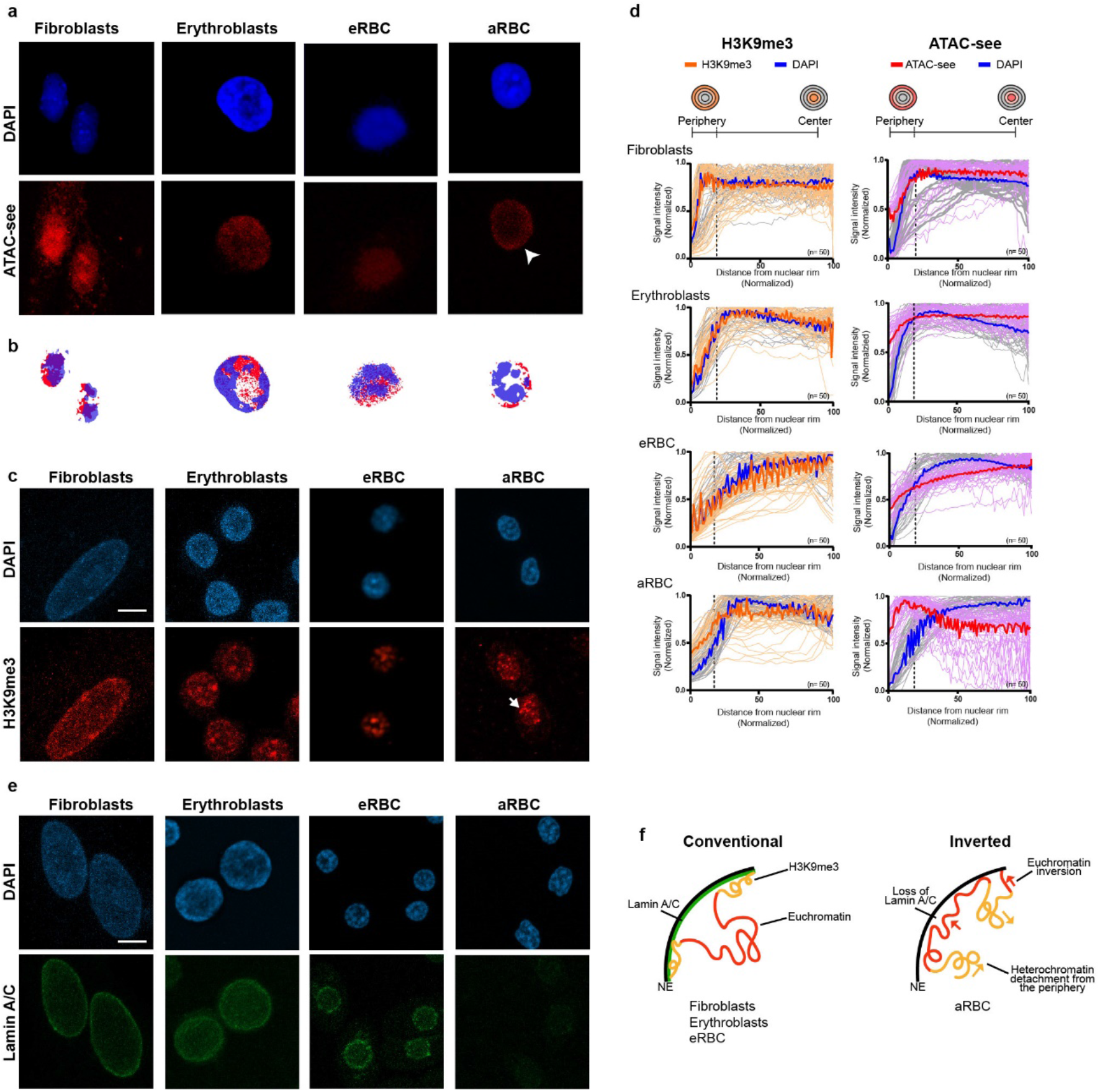
Chromatin remodeling during chicken erythropoiesis results in an inverted nuclear architecture. **a**, Imaging of open chromatin by ATAC-see (red) and DAPI (blue, nuclei) in fibroblasts and erythroid cells. Repositioning of the accessible chromatin towards the nuclear periphery in aRBC is indicated with an arrowhead. **b**, Brightest pixels enriched in ATAC-see (red) or DAPI (blue). **c**, Immunofluorescence imaging of H3K9me3 (red) and DAPI. In aRBC the heterochromatin is relocated towards the nuclear center (arrows). **d**, For each cell type, H3K9me3, ATAC-see and DAPI intensity is presented as a function of distance to the nuclear periphery. Individual tracks for 50 nuclei are presented in lighter colors and the mean intensity is presented in bold. **e**, Fluorescence imaging of Lamin A/C proteins (green) and DAPI. In every case, maximal projections of three-dimensional (3D) stacks are presented. Scale bar = 5 µm. **f**, In contrast to the conventional nuclear architecture in fibroblast, erythroblasts, and eRBC in aRBC the loss of the nuclear lamin A/C leads to heterochromatin detachment from the nuclear periphery and euchromatin inversion.

ATAC-see of aRBC revealed a unique nuclear organization in which euchromatin is located predominantly at the periphery and less abundant at the center of the nucleus (Fig. 1a; arrowhead) as opposed to the conventional nuclear architecture observed in fibroblasts, erythroblasts, and eRBC. Next, we applied a Gaussian filter to the raw images using a threshold to obtain all pixels brighter than the mean in both ATAC-see and DAPI channels in each nucleus (see methods). The images generated confirmed that euchromatin is enriched at the nuclear periphery in aRBC while the DAPI signal is more intense at the nuclear center (Fig. 1b; Extended data 1c).

Heterochromatin is also relocated in chicken erythrocytes as shown by H3K9me3 immunostaining (Fig. 1c; arrow). In eRBC and aRBC H3K9me3 is preferentially segregated into foci located at the center of the nucleus instead of the conventional positioning of constitutive heterochromatin at the nuclear envelope (NE) seen in fibroblasts and erythroblasts.

We validated the inverted organization of aRBC nuclei by plotting the normalized mean intensity of H3K9me3 and ATAC-see as a function of distance from the nuclear periphery (Fig. 1d). In fibroblasts and erythroblasts, H3K9me3 signal gradually decreases from the NE to the interior, where ATAC-see signal is higher. Consistently with a conventional nuclear architecture. In contrast, in eRBC and aRBC H3K9me3 increases from the nuclear periphery to the interior reflecting the position of the heterochromatic foci we observed in these cells. In agreement with the inversion of euchromatin observed in aRBC, ATAC-see signal intensity in these cells is higher at the nuclear periphery and decreases to the nuclear interior. Pearson’s correlation analysis between signal intensity and distance from the nuclear periphery also recapitulates the inversion of euchromatin in aRBC (Extended data 1d). Thus, terminal erythroid differentiation results in repositioning of heterochromatin away from the NE in eRBC and aRBC, and the relocation of euchromatin towards the periphery in aRBC.

The unique inverted nuclear architecture observed in aRBC has been previously shown to occur in the rod photoreceptor cells of nocturnal mammals in response to downregulation of the Lamin B receptor (LBR) and absence of Lamin A/C expression ^32^. We asked whether the chromatin inversion observed in aRBC could be explained by changes in the NE composition. Like the nuclear inversion in rod cells, Lamin A/C is present in the cells with conventional nuclear architecture but lost in the aRBC nuclei (Fig. 1e). Lamin B1 was detected in all cells studied (Extended data 1e).

Taken together, these results show a complex reorganization of the nuclear architecture in chicken erythrocytes. Cells with a conventional nuclear architecture locate heterochromatin at the NE while euchromatin is preferentially positioned at the nuclear interior. During terminal differentiation, heterochromatin moves away from the NE at foci and loss of Lamin A/C in aRBC allows repositioning of euchromatin towards the nuclear periphery in an inverted architecture (Fig. 1f).

### Chicken erythrocytes show a global downregulation of gene expression except for small RNAs and erythroid-related genes

We wondered if the accessible chromatin detected by ATAC-see in the chicken erythrocytes reflected sites of ongoing transcription in these cells. Quantitation of total RNA extracted from erythrocytes showed a dramatic decrease in RNA content in eRBC and aRBC (Fig. 2a) in agreement with a previous report of low levels of RNA synthesis in eRBC measured by radioactivity incorporation^10^. Furthermore, the electrophoretic profile of the RNA obtained shows changes in the diversity of RNA species in eRBC and aRBC with a gradual loss of the rRNA bands while small RNA molecules are still detectable (Fig. 2b).

**Fig. 2.**
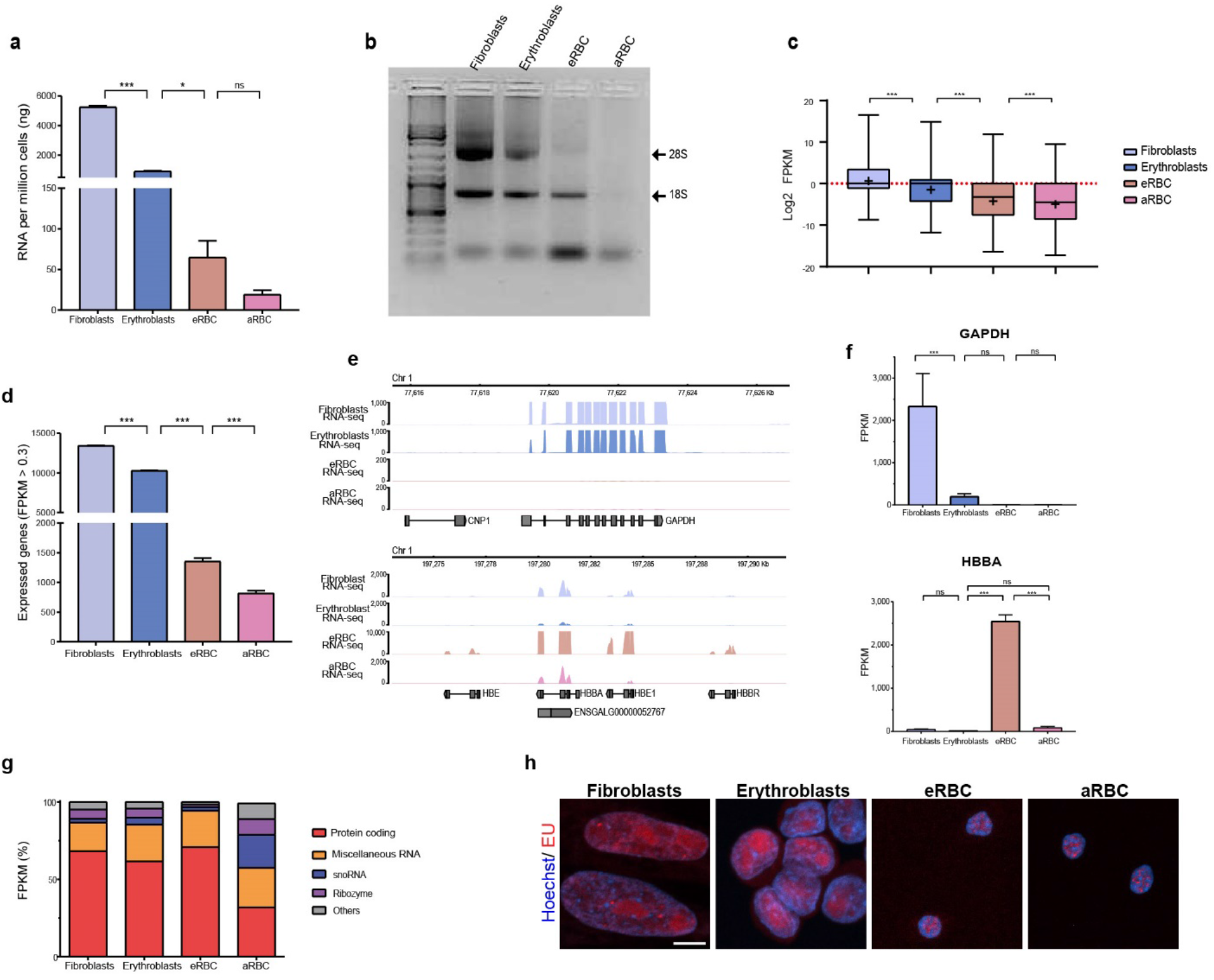
Chicken erythrocytes show a dramatic decrease in transcription but continue expressing small RNAs and erythroid genes. **a**, Quantification of total RNA obtained from chicken fibroblasts, erythroblasts, eRBC, and aRBC. Shown are values and means of 3 independent replicates. Asterisks indicate P values based on one-way ANOVA followed by Fisher’s LSD post-hoc test. *P < 0.05; ***P < 0.001; ns not significant differences. **b**, Representative picture of an agarose gel electrophoresis of total RNA. **c**, Mean Log2 FPKM values of three RNA-seq experiments. Mean is shown as +. Statistical differences were determined based on a one-way ANOVA followed by Dunn’s post hoc test. *** P < 0.01. **d**, Number of expressed genes (FPKM > 0.3) according to RNA-seq experiments. Asterisk indicate statistical differences based on one-way ANOVA followed by Tukey’s post hoc test. *** P <0.01 **e**, RNA-seq tracks for GAPDH (top) and the beta-globin locus (bottom). **f**, FPKM values for GAPDH (top) and HBBA (bottom) genes, shown are the values of three replicates with standard deviation. Asterisk indicate statistical differences based on one-way ANOVA followed by Tukey’s post hoc test. ** P < 0.05; *** P <0.01; ns not significant differences. **g**, RNA diversity expressed as the percentage of total FPKM that correspond to protein coding genes, Miscellaneous RNA, snoRNA, ribozyme or others. **h**, Fluorescent labeling of nascent RNA with 5-ethynil uridine (EU, red). Maximal projections are presented. Scale bar = 5 µm

To further characterize the transcriptional state of chicken erythrocytes we performed total RNA-seq experiments. A global downregulation of gene expression in the erythrocytes was evidenced by a marked decrease in FPKM values compared to other cell types (Fig. 2c). We identified all genes that were expressed (FPKM > 0.3) and found a small set of genes that remain active in the erythrocytes (1,354 in eRBC and 804 in aRBC) in agreement with RT-qPCR experiments (Fig. 2d-f; Extended data 2a-d). Of note, most FPKMs come from small RNA molecules such as snoRNAs and miscellaneous RNAs in eRBC and aRBC, consistent with the electrophoretic pattern of erythroid RNA extractions (Fig. 2g).

Differential expression analysis was conducted between the different erythroid stages. We found 810 and 428 differentially expressed genes between eRBC or aRBC and erythroblasts respectively (Extended data 2e). Gene ontology analysis of upregulated DEGs showed that chicken erythrocytes exhibit upregulation of genes related to oxygen-carrier activity, erythroid differentiation, and immune activity (Extended data 2f). The presence of active transcription in chicken erythrocytes was corroborated *in vivo* by incorporation of 5-ethynyl uridine (EU) to nascent RNA and subsequent detection and imaging of EU (Fig. 2h). Hence, chicken erythrocytes show a global downregulation of gene expression but retain transcription of a small number of genes associated with erythroid function.

Nascent RNA labelling allowed us to study the spatial location of transcription. As expected, RNA synthesis in fibroblasts and erythroblasts was distributed throughout the nucleus but more prominent at the center (Fig. 2h). In contrast, transcription in the erythrocytes was spatially restricted into foci, indicating a high degree of compartmentalization inside the nucleus. However, the presence of open chromatin at the NE cannot be explained by active transcription since transcriptional foci were not preferentially located at the nuclear periphery in aRBC.

### The paused RNA polymerase retains accessibility at silent promoters in chicken erythrocytes

We performed ATAC-seq experiments to identify the genomic regions that remain accessible in the chicken erythrocytes. We verified that ATAC-seq replicates of each cell type were highly correlated (*R* > 0.92; Extended data 3a). Overall, we identified fewer ATAC-seq peaks in erythroid cells compared to fibroblasts, suggesting that chromatin accessibility is already diminished at the erythroblast stage (Fig. 3a). In fact, we identified similar numbers of ATAC-seq peaks in erythroblasts, eRBC, and aRBC regardless of the genome-wide decrease in gene expression in the erythrocytes. Most of the peaks in the erythrocytes represent constitutively open regions or shared with the erythroblasts, indicating that these regions retain their accessibility throughout terminal erythroid differentiation (Fig. 3b).

**Fig. 3.**
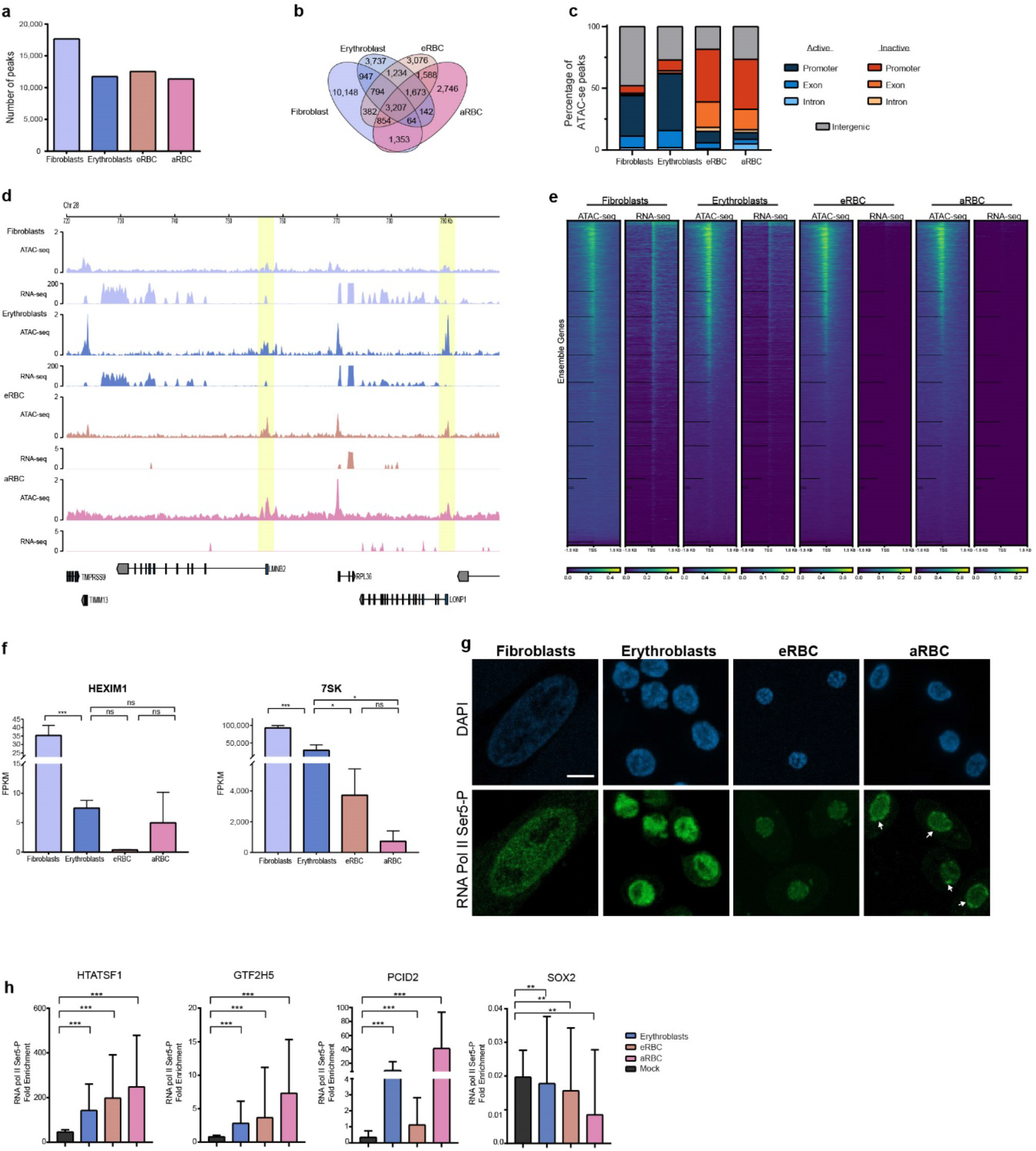
Open chromatin in the chicken erythrocytes comprises paused promoters of silenced genes. **a**, Number of ATAC-seq peaks detected in each cell type. **b**, Venn diagram of ATAC-seq peaks in each cell type. **c**, Annotation of ATAC-seq peaks at intergenic regions, promoters, exons and introns of transcriptionally active or inactive genes. **d**, ATAC-seq and RNA-seq tracks showing an example of inactive genes in eRBC and aRBC that retain accessibility at their promoter (yellow) in absence of transcription. **e**, ATAC-seq and RNA-seq CPM ±1.5 Kb of annotated TSSs. **f**, Expression levels of HEXIM1 and 7SK, part of the 7SK snRNP involved in RNA polymerase pausing (n = 3). Asterisk indicated significant differences based on a one-way ANOVA followed by Tukey’s post hoc test. **g**, Imaging of the RNA pol II Ser5-P. In aRBC the RNA pol II Ser5-P accumulates at the nuclear periphery (arrow) Maximal projections are presented. Scale bar = 5 µm. **h**, ChIP-qPCR against RNA pol II Ser5-P. ATAC-seq peaks at the promoter of HTATSF1, GTF2H5 and PCID2 were analyzed. Fold enrichment was calculated against amplification in the promoter of SOX2. SOX2 enrichment was calculated against input. Mean and standard deviation from 4 replicates is presented. Statistical differences between against a mock control were calculated with a t-test.

We classified the ATAC-seq peaks according to their genomic location and transcriptional state and found that many peaks in eRBC and aRBC correspond to silenced genes that remain accessible at their promoters (Fig. 3c-e; Extended data 3b). Promoter-proximal pausing of the RNA polymerase II has been shown to maintain promoter chromatin accessibility and the gene’s potential to be activated^33^. We hypothesized that RNA pol II pausing could be responsible for preserving the accessible promoters in the genome of eRBC and aRBC.

RNA pol II pausing is promoted by the inhibition of the positive elongation factor p-TEFB by the 7SK small nucleoprotein complex formed by the snRNA 7SK, HEXIM1/2, LARP1, and MePCE^34,35^. Interestingly, 7SK and HEXIM1 are highly expressed in the erythrocytes (Fig. 3f). Furthermore, immunofluorescence of the RNA pol II engaged in transcriptional initiation (Ser5-P) was readily detected in all cell types with no significant changes in signal intensity (Fig. 3g; Extended data 3d), whereas the signal from the elongating CTD isoform (Ser2-P) is diminished in eRBC and aRBC (Extended data 3c, e). In addition, we found that RNA pol II Ser5-P is spatially located at the nuclear periphery in aRBC (Fig 3g; arrows) following the same distribution as the euchromatin detected by ATAC-see (Extended data 3f-g). Finally, we confirmed the presence of RNA pol II Ser-5P in the open promoter regions of genes without expression in the erythrocytes by ChIP-qPCR (Fig. 3h; Extended data 3h). In contrast, no enrichment of RNA pol II Ser5-P was detected on the inaccessible *Sox2* gene promoter. These results indicate that despite global reduction of gene expression, chicken erythrocytes retain expression of genes involved in RNA pol II pausing and accessibility at gene promoters due to the presence of RNA pol II in a promoter-proximal paused state.

### The genome of eRBC and aRBC is hyper-compartmentalized and structured in mini domains at open chromatin regions

Recent Hi-C experiments conducted in aRBC have evidenced a global rearrangement in their 3D genome organization characterized by an increase in long-range interactions and absence of Topologically Associated Domains (TADs)^36^. However, characterization of the chromosome organization at distinct stages of erythroid differentiation is still lacking. We sought to explore how the observed changes in nuclear architecture, chromatin accessibility, and gene expression relate to 3D genome organization and generated Hi-C contact maps for chicken fibroblasts, erythroblasts, eRBC, and aRBC (Extended data 4a).

High-order chromatin organization is defined by the segregation of genomic regions into A and B compartments corresponding to active and inactive chromatin, respectively^16^. These compartments are observed in whole-chromosome Hi-C contact maps as a plaid pattern of long-range interactions. Visual inspection of Hi-C maps shows a more pronounced plaid pattern in eRBC and aRBC than in erythroblasts (Fig. 4a-b), indicative of the strengthening of chromatin segregation into A and B compartments in these cells. Comparison of Hi-C maps also shows a decrease in interactions along the diagonal. Accordingly, we detected an increase in long-range (> 10 Mb) interactions in the erythrocytes while short-range interactions are less frequent (Fig. 4c). These results indicate that the genome of chicken erythrocytes is strongly compartmentalized, as evidenced by the presence of well-defined heterochromatin and transcriptional foci in these cells.

**Fig. 4.**
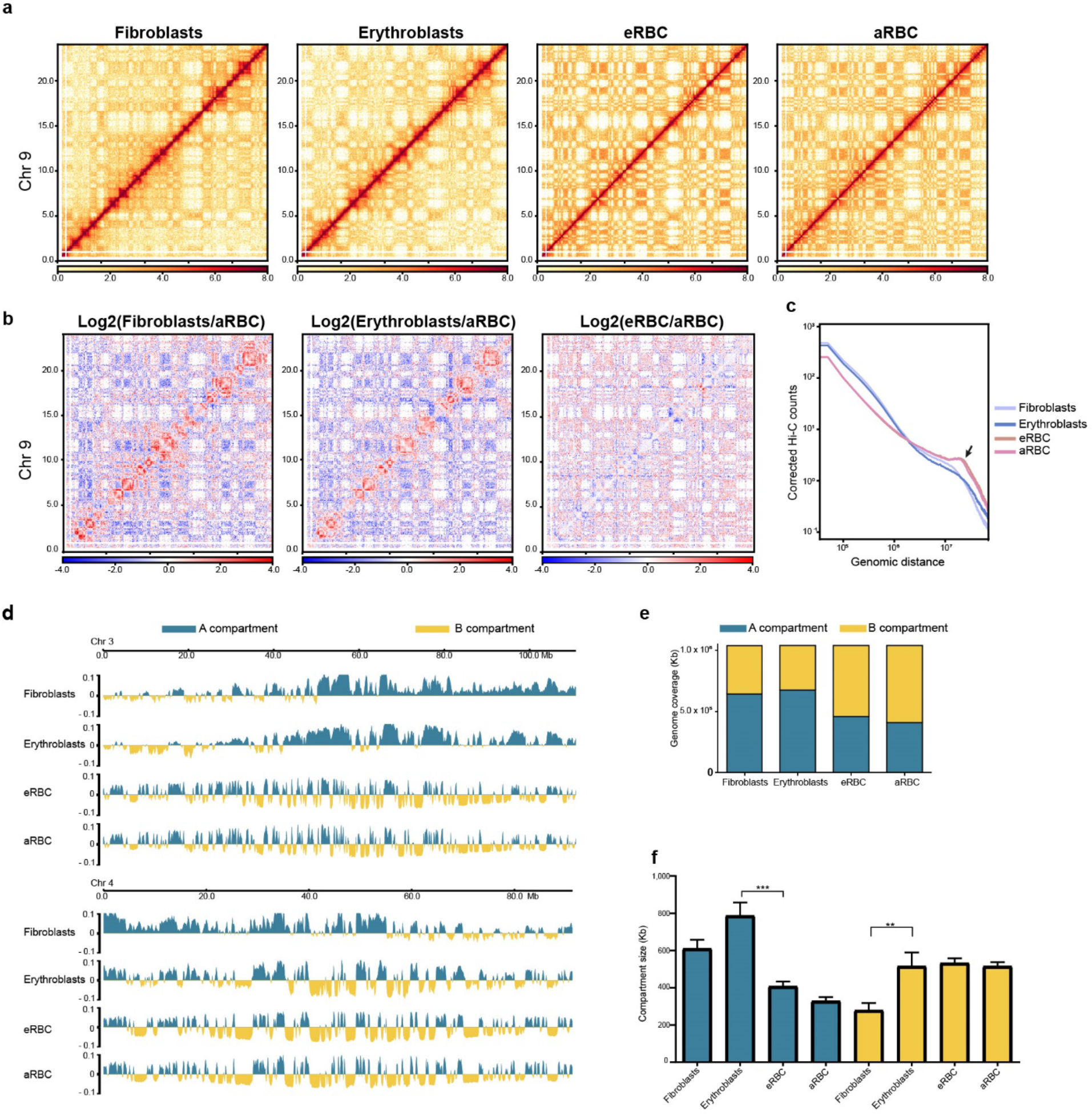
Chicken erythrocytes have hyper-compartmentalized genomes. **a**, Hi-C contact matrices of chromosome 9 (Mb 0 -24). **b**, Hi-C contact matrices showing the log2 fold change between each cell type and aRBC. **c**, Hi-C contacts counts as a function of genomic distance. An increase in long-range interaction in eRBC and aRBC is indicated with an arrow. **d**, A (blue) and B (yellow) compartment identification by PCA analysis in fibroblast, erythroblasts, eRBC, and aRBC in chromosome 3 (top) and 4 (bottom). **e**, Genome coverage of the A and B compartment. **f**, Average size of A and B compartment regions. Asterisks indicate P values based on one-way ANOVA followed by Bonferroni’s post-hoc test. **P < 0.01; ***P < 0.001.

Identification of A and B compartments showed a hyper-compartmentalization pattern in the genome of the erythrocytes compared to erythroblasts, characterized by the disruption of long A-type regions by the emergence of regions in the B compartment (Fig. 4d). In fact, most of the genome falls into the B compartment in the erythrocytes, as regions in the A compartment are reduced in size compared to erythroblasts and fibroblasts (Fig. 4e-f). This hyper-compartmentalized pattern with the B compartment “invading” the A compartment in the erythrocytes reflects the genome-wide reduction in transcription in these cells.

Strong compartmentalization and decrease in short-range interactions make visual identification of topological domains difficult in eRBC and aRBC (Fig. 5a) and it has been suggested that chicken erythrocytes lack TADs^36^. TAD calling using insulation index^37^ uncovered a profound loss of TAD structures in chicken erythrocytes except at regions where a closer inspection reveals a local enrichment of short-range interactions (Fig. 5b;Extended data 4b). These domains are fewer and smaller (with an average size of 241 Kb) than TADs observed in erythroblasts and fibroblasts (Fig. 5c-d), hence we called these structures min-domains. Of note, only a small fraction of the genome remains structured in mini domains (24.18% in eRBC and 20.4% in aRBC) (Fig. 5e).

**Fig. 5.**
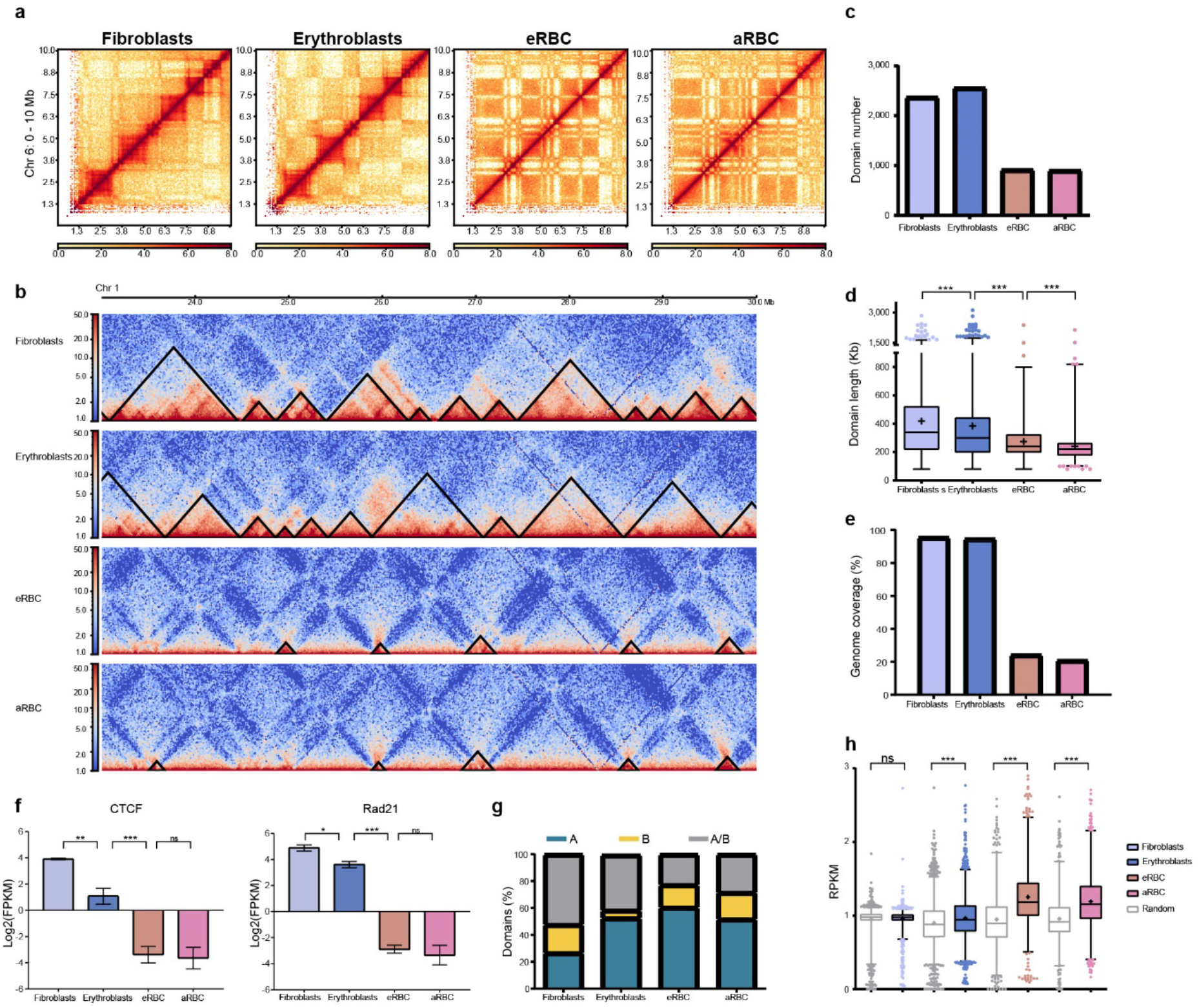
Chicken erythrocytes present global TAD loss but retain structure at mini domains enriched in open chromatin. **a**, Hi-C contact matrices of the first 10 Mb of chromosome 6. TADs are clearly visible in fibroblast and erythroblasts while the most prominent signal in erythrocytes corresponds to the plaid pattern of compartments. **b**, Domains identified in each cell type indicated by black triangles. Number (**c**,) and average size of the domains is presented (**d**,). **e**, Percentage of the genome structured in domains. **f**, Expression levels of architectural proteins based on RNA-seq data. Asterisks indicate P values based on one-way ANOVA followed by Tukey’s post hoc test. ns = P > 0.05; * P < 0.05, ** P < 0.01, ***P < 0.001. **g**, Percentage of domains that contain only A compartment regions (blue), B compartment regions (yellow), or both (gray). **h**, ATAC-seq RPKM count inside domains or a random control. Asterisks indicate P values based on one-way ANOVA followed by Kruskal-Wallis post hoc test. ns = P > 0.05; ***P < 0.001.

Topological domains insulation depends on the presence of the architectural proteins CTCF and cohesin at the domain boundaries^24,38,39^ and auxin-inducible depletion of CTCF or the cohesin-loading factor Nipbl causes genome-wide loss of topological domains^22,23^. Interestingly, CTCF and Rad21 are downregulated in eRBC and aRBC (Fig. 5f). Thus, downregulation of architectural proteins might explain the loss of topological domains in chicken erythrocytes.

Notably, mini domains demarcate regions in the A compartment and are enriched in ATAC-seq reads (Fig. 5g-h). Taken together, these results indicate that despite a global loss of TADs in the genome of eRBC and aRBC, euchromatic regions remain structured in mini domains.

### Chicken RBC genomes are organized around paused promoters

We hypothesized that the presence of local structure at mini domains reflected transcriptional activity inside erythroid mini domains. Indeed, mini domains are enriched in RNA-seq reads (Extended data 4c). However, visualization of Hi-C maps together with ATAC-seq and RNA-seq tracks showed that mini domains are often found around open regions of the genome that lack transcription (Fig. 6a; Extended data 4b).

**Fig. 6.**
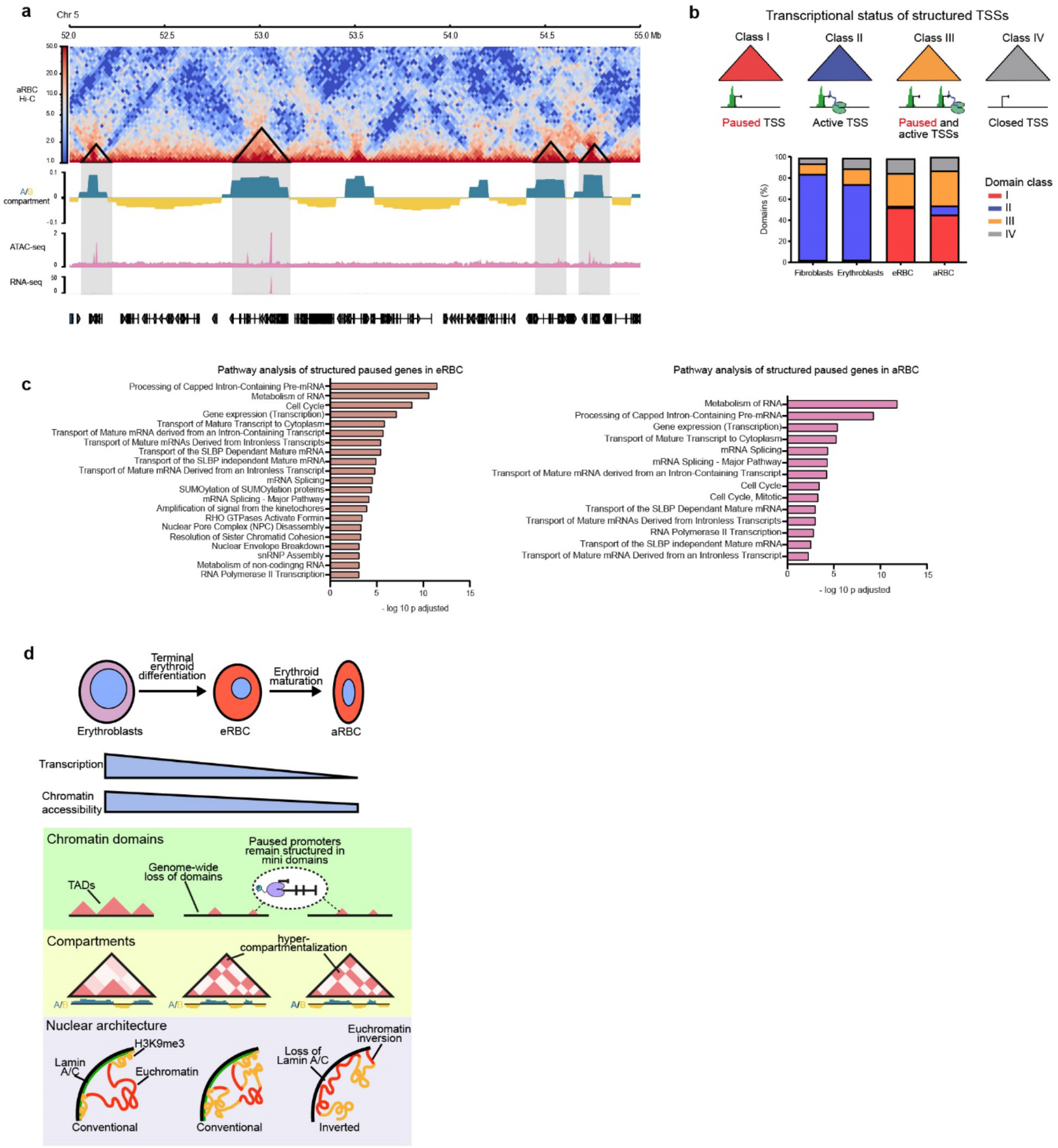
Mini domains contain and organize paused promoters in chicken erythrocytes. **a**, Hi-C contact matrix from aRBC (5: 52 – 55 Mb), the mini domains identified (black triangles) and the corresponding compartment signal, ATAC-seq, and RNA-seq tracks. aRBC mini domains contain ATAC-seq peaks independent from transcription (grey). **b**, Distribution of mini domains according to the transcriptional status of the contained TSSs into Class I (paused TSSs), Class II (active TSSs), Class III (Both paused and active TSSs) or Class IV (closed TSSs). **c**, Gene Ontology terms of paused TSSs structured in mini domains. **d**, Terminal erythroid differentiation in chicken results in a dramatic drop in transcription and chromatin accessibility that is exacerbated during erythroid maturation. Erythrocyte genomes lose TADs genome-wide but retain mini domains structured around open paused promoters with presence of the stalled RNA polymerase. At the compartment level the erythrocyte genome is hypercompartmentalized and rich in long-range interactions. Lastly, in aRBC the absence of Lamin A/C causes chromatin inversion.

We classified topological domains by the transcriptional and chromatinic state of their TSSs (Fig. 6b). Domains within the class I contain open promoters of genes without expression, herein referred to as “paused” promoters while TSSs within class II domains are open and transcriptionally active. Class III domains contain both paused and active promoters and class IV are depleted of open TSSs. This classification evidenced that the presence of paused promoters is a common feature in mini domains as more than 80% of mini domains fall into either class I o III.

Interestingly, paused genes that reside within mini domains in eRBC (3,458 genes) and aRBC (2,804 genes) are involved in regulation of gene expression, RNA processing and transport of mature RNAs to the cytoplasm (Fig. 6c). Examples of such paused genes are the transcriptional elongation factors *Htatsf1* and *Gtf2h5*^40,41^, and the nuclear export protein *Pcid2*^42^ (Fig. 3h). Paused genes in erythroblasts (726 genes) and fibroblasts (791 genes) were not enriched in any pathway. One possible explanation for this observation is that the preserved mini domains together with the paused RNA pol II bookmark the necessary components of the transcriptional machinery and RNA processing complexes to rapidly resume gene expression in response to environmental cues. For example, transcriptional activation of the erythroid genome has been observed in the nucleated erythrocytes of fish and birds in response to immune challenges^43^. In summary, we show that paused promoters of genes involved in gene expression and RNA metabolism remain organized into mini domains despite a global loss of TADs in chicken eRBC and aRBC.

## Discussion

Here we analyzed thoroughly the changes in chromatin accessibility and organization that accompany terminal erythroid differentiation and downregulation of gene expression in the nucleated chicken erythrocytes. We found that loss of Lamin A/C results in a complete inversion of the nuclear architecture in aRBC. Furthermore, the erythrocyte nucleus is a highly compartmentalized organelle that confines euchromatin, transcription and heterochromatin into readily observable distinct foci. Downregulation of architectural proteins greatly affects TADs genome-wide. However, RNA pol II pausing retains accessibility at open promoters and supports short-range chromatin interactions, thus forming mini domains around genes important in RNA synthesis and RNA metabolism. Our results uncover a complex reorganization process at different genomic scales with important implications in the physiology of nucleated erythrocytes and the mechanisms that govern genome organization.

Chromatin condensation and gene silencing are common traits of mammalian and non-mammalian erythropoiesis^5,9,44^. Nucleated erythrocytes are commonly described as transcriptionally inert cells^12^ and defects in heterochromatin stability or gene silencing are considered hallmarks of some myelodysplastic syndromes and anemia in mammals and birds^45-47^. Here, we provide evidence that both eRBC and aRBC actively transcribe a reduced number of genes (1,354 in eRBC, 804 in aRBC) including hemoglobin and genes involved in gas exchange and transport. These results challenge the notion of nucleated erythrocytes as transcriptionally inactive cells and suggest a functional role of the nucleus in the erythroid physiology.

By direct imaging of RNA synthesis in chicken erythrocytes we found that transcription is confined into discrete active foci. In fact, segregation of euchromatin and heterochromatin into different compartments is exacerbated as evidenced by imaging and Hi-C data. Chromatin compartmentalization prevents abnormal contacts between euchromatic and heterochromatic regions that may result in heterochromatin spreading and ectopic gene repression^48^. The confinement of transcription into foci in erythrocytes probably reflects the requirement of bringing together the remaining active regions of the genome to favor transcription in an environment where transcriptional factors are scarce.

Heterochromatin is also relocated in eRBC and aRBC. In aRBC, loss of Lamin A/C leads to a complete inversion of nuclear architecture in which heterochromatin detaches from the nuclear envelope and moves towards the center of the nucleus. The atypical spatial organization of euchromatin at the nuclear rim was described in the rod photoreceptors of nocturnal mammals where loss of heterochromatin tethering to the nuclear envelope is triggered by downregulation of LBR and absence of Lamin A/C^32^.

In recent years, heterochromatin has emerged as a key factor mediating genome organization and heterochromatin repositioning has important effects on cellular function ^49^. For example, the inverted nuclear architecture in rod cells is thought to maximize light transmission in the retina and disposition of accessible chromatin at the nuclear rim in neutrophiles prepares the cells for initiation of NETosis^31^. Moreover, heterochromatin detachment and relocation cause gene misexpression in laminopathies and senescence^49^. An increase in heterochromatin-mediated long-range interactions is observed in the last nucleated stage of human erythropoiesis^50^ and Lamin B cleavage by caspase-3 and transient nuclear opening has been shown to promote chromatin condensation in the erythroblasts of mouse and human^4,45^. Together with our results, these evidences suggests that nuclear envelope remodeling, compartment strengthening, and heterochromatin relocation may be a conserved mechanism to ensure proper chromatin compaction.

The molecular forces driving genome compartmentalization are not entirely understood. Interestingly, knock-out of the cohesin loading complex Nipbl or the cohesin subunit Rad21 causes a general enhancement of compartmentalization^23,51^.Thus, compartmentalization may be regulated by the binding of cohesin to chromatin and loop-extrusion. Here we found Rad21 to be downregulated in eRBC and aRBC which provides a possible explanation for the hyper compartmentalized state of their genomes.

Recent Hi-C experiments in chicken aRBC and the immature polychromatic erythrocyte using a six-cutter restriction enzyme showed a genome-wide depletion of TADs in these cells^36^. Although strong compartment signal obscures the presence of chromatin domains in the Hi-C matrices, here we provide compelling evidence of local interacting mini domains that persist structured in the terminal differentiated eRBC and aRBC. TADs are formed when the loop-extrusion activity of the cohesin complex is blocked by the presence of CTCF bound at convergently oriented binding-sites located at TAD boundaries^52^. Ablation of CTCF results in a global loss of TADs except at regions where low levels of CTCF are able to maintain TAD boundaries^22,53^. Similarly, degradation of Rad21 and Nipbl eliminates chromatin domains^23,51^. Here we show that transcription of CTCF and Rad21 decreases in the nucleated erythrocytes of chicken, which may account for a loss of chromatin domains. TAD disturbance has been recently shown to occur in human erythroblasts without evident changes in chromatin compartmentalization^50^. Thus, global genome reorganization represents a conserved feature of terminal erythropoiesis that reflects the particular transcriptional state of these cells.

The contribution of RNA polymerase to the establishment and maintenance of genome organization is still a matter of debate. The emergence of TADs during development coincides with the transcriptional activation of the zygotic genome^28^. Inhibition of transcriptional elongation weakens TAD insulation but does not preclude TAD emergence^19,28,54^. Hence, transcription is not necessary for the establishment of TADs but might play a role in maintaining chromatin organization by other associated processes like chromatin opening, the assembly of the pre-initiation complex, and recruitment of transcription factors and the transcriptional machinery^28^. Moreover, RNA pol II dictates cohesin loading into chromatin and rapid degradation of RNAPII prevents the reestablishment of chromatin folding in cells following exit from mitosis^55^. Our results show that mini domains in the chicken erythrocytes frequently contain paused promoters occupied by the RNA pol II-Ser5, suggesting that RNA Pol II binding is enough to maintain local organization and insulation of small domains.

Promoter-proximal pausing of RNA pol II has been shown to serve as a marker that primes silenced genes to be rapidly activated during development and in response to environmental cues^56^. Here we present evidence that suggest that RNA pol II pausing represents a key mechanism by which chicken nucleated erythrocytes maintain low levels of transcription. We show that eRBC and aRBC sustain elevated levels of transcription of components of the snRNP 7SK complex involved in polymerase pausing and observed the presence of RNA pol II Ser5-P in the nucleus of these cells despite a global decrease in transcription. In agreement with early run-on experiments that identified the presence of a stalled form of the RNA pol II in the beta-globin genes in hen erythrocytes^57^. A recent study suggests that regulation of RNA pol II activity is important during terminal erythroid maturation in humans^58^.

Finally, paused promoters in eRBC and aRBC are enriched in categories concerning transcriptional elongation, RNA processing and nucleocytoplasmic transport of mature RNAs. We hypothesize that Pol II pausing prepares chicken nucleated erythrocytes to restore the expression of components of the transcriptional and RNA processing machineries to mount an efficient response to stimulus like immune challenges^43,59^.

Our results reveal a dramatic reorganization of the genome at different scales during the formation of nucleated erythrocytes in chicken including the inversion of euchromatin regions to the nuclear periphery in aRBC and highlight the role of RNA pol II in the maintenance of insulated domains in this highly repressive chromatin environment. We show that nucleated erythrocytes maintain a collection of accessible promoters by promoter-proximal pausing of the RNA pol II at genes important to mount a transcriptional response.

## Methods

### Blood collection and cell culture

All applied procedures were approved considering the animal welfare practices according to the IFC animal facility board. Blood of 10-day old chicken embryos or 28-38 weeks-old White-leghorn hens was collected in cold phosphate saline buffer with 50 mM EDTA. Cells were centrifuged at 1,000 rpm, 4°C for 10 minutes and washed three times and counted on a Neubauer chamber. Chicken fibroblasts were prepared from 10-day old chicken embryos

Chicken fibroblasts and the erythroblasts HD3 cell line were cultured in Dulbecco’s modified Eagle’s medium (DMEM; GIBCO) supplemented with 10% fetal bovine serum (Biowest), 2% chicken serum and penicillin/streptomycin (Invitrogen).

### ATAC-seq library preparation

ATAC-seq experiments were performed as previously described^60^ with minor modification for chicken cells. 150,00 cells lysed on lysis buffer (10 mM Tris-Cl pH 7.4, 10 mM NaCl, 3 mM MgCl2, 0.01% Igepal) for 15 minutes on ice. Fibroblasts were sieved through a 70 µm cell-strainer and dounce homogenized before lysis. Nuclei were centrifuged at 600 x g, 4°C for 10 minutes and washed with 50 µL of lysis buffer. Nuclei were incubated with the Tn5 transposase (Illumina) and tagmentation buffer (Illumina) for exactly 30 minutes at 37°C. After tagmentation, fragments were purified using MinElute columns (Qiagen). ATAC-seq libraries were amplified by 10 PCR cycles using the Nextera index system (Illumina). Libraries were quantified in a Qubit system (ThermoFisher) and fragment size was determined on a TapeStation 4200 system (Agilent). Paired-end sequencing of biological replicates was conducted in a HiSeq2500, NextSeq or NovaSeq machine.

### ATAC-seq data processing

ATAC-seq reads were aligned against the chicken genome (GRCg6a) using Bowtie2^61^ with the default parameters for paired-end reads. BAM files were generated using samtools (samtools view), sorted (samtools sort), and indexed (samtools index). Duplicates (samtools markdup) and reads longer than 150 bp (samtools view) were removed. Signal tracks were obtained using deeptools (bamCoverage –normalizeUsing CPM) and visualized with pyGenomeTracks^62^.

Accessible peaks were identified on each replicate using MACS2^63^ callpeak function with options -f=BAMPE and -q 0.01. Consensus peaks conserved in between the two biological replicates were conserved for subsequent analysis.

### RNA purification and RNA-seq library preparation

Total RNA was extracted using TRIzol following the manufacturer’s instructions and the integrity of the material was evaluated by electrophoresis. RNA-seq libraries were prepared starting from 1µg of RNA using the TruSeq Stranded Total RNA Ribo-Zero H/M/R Gold kit (Illumina). Libraries were quantified in a Qubit system (ThermoFisher) and fragment size was determined on a TapeStation 4200 system (Agilent). Three biological replicates per cell type were sequenced on a HiSeq2500 machine, paired end 50 bp.

### RNA-seq data processing

RNA-seq data was analyzed using the SnakePipes pipeline^64^. Briefly, RNA-seq reads were mapped to the GRCg6a genome using STAR^65^, raw counts were obtained using featureCounts^66^, and FPKM were calculated. Because RNA content in chicken erythrocytes is significantly lower than in the other cell types, raw counts and FPKM were normalized for the number of cells required to obtain 1 µg of RNA. Genes with ≥ 0.3 FPKM were considered as expressed genes. Differential expression analysis was performed with DESeq2^67^ using an FDR value of 0.5 and FC of 2.

### Gene Ontology analysis

Gene ontology and pathway enrichment analysis were done using g:Profiler^68^ using default parameters. All significant biological processes and pathways have a p value < 0.01.

### Hi-C library preparation

Hi–C libraries were generated as previously described^69^. Briefly, 10 × 10^6^ cells were fixed with formaldehyde 2% for 10 minutes at room temperature. The genomes were digested with MboI (NEB) overnight and 5′ overhangs were filled with Biotin-14-dATP (Invitrogen) and religated in nuclei. DNA fragments were purified and shared to an average size of 500 bp using a Covaris machine. The sheared DNA was end-repaired, adenine tailed, and size-selected. Biotinylated fragments were pulled down using MyOne Streptavidin C1 DynaBeads (Invitrogen) and ligated to Truseq adapters (Illumina). Hi-C libraries were amplified using 5-6 PCR amplification cycles. Libraries were quantified in a Qubit system (ThermoFisher) and fragment size was determined on a TapeStation 4200 system (Agilent). Two biological replicates per cell type were paired end sequenced using a HiSeq2500 or NextSeq machine.

### Hi-C data processing

Mapping, filtering, correction, and generation of Hi-C matrices was done using HiC-Pro^70^. Briefly, read pairs were mapped against the chicken genome (GRCg6a) using Bowtie2 with HiC-Pro default parameters. Valid pairs were used to generate raw and ICE normalized matrices at 20 Kb, 50 Kb and 100 Kb bin resolution. Correlation between Hi-C replicates and counts vs distance plots were generated with HiCExplorer^62^ with a bin size of 50 Kb and replicates were merged (samtools merge). Matrices were normalized to the smallest number of valid pairs to correct for differences in sequencing depth. Heatmaps of normalized contact matrices and matrix comparisons were performed with HiCPlotter^71^.

### TAD calling

Dense Hi-C matrices generated with HiC-Pro (sparseToDense.py) and TAD calling was conducted with TADtool^37^ using the insulation score algorithm at a 20 Kb resolution with a window size of 100 Kb. TADs identified showed good agreement with visual inspection of Hi-C matrices.

### Compartment analysis

Chromatin compartments were identified using HiCExplorer (hicPCA) at 50 Kb resolution. RNA-seq tracks were used to assign the sign of PCA results.

### Antibodies

The following antibodies were used for immunofluorescence experiments: anti-H3K9me3 (Millipore, #07-442), anti-RNA pol II Ser5-P (Santa Cruz, #SC-47701), anti-RNA pol II Ser2-P (Abcam, #ab5095), Lamin B1 (Abcam, #ab16048), anti-Lamin A/C (Santa Cruz, #SC-376248), anti-rabbit Alexa Fluor 594-conjugated (Life Technologies, #R37117), and anti-mouse FITC-conjugated (Millipore, AP192F).

### Immunofluorescence

Cells were plated and cultured on coverslips pretreated with Poly-L-lysine (Electron Microscopy Sciences), then fixed with formaldehyde 1% and permeabilized with Igepal 0.1%. Fixed cells were washed with PBS and preincubated with 5% bovine serum albumin (BSA) for 1 hour and then incubated with primary antibodies in PBS with 0.2 BSA overnight at 4°C. After three washes with PBS, cells were incubated with the secondary antibody for 1 hour and stained with DAPI for 20 minutes, then mounted using Fluoromount (Sigma). Image acquisition was performed in a Zeiss LSM800 confocal microscope.

### ChIP-qPCR

For ChIP-qPCR experiments, four biological replicates per cell type were fixed in 1% formaldehyde for 10 min. Samples were sonicated for 10 cycles of 30sec on/60seg off with 30% amplitude using a Cole-Parmer 500-watt ultrasonic Processor. Chromatin immunoprecipitation was performed using 5µg of anti-RNA pol II Ser5-P antibody (Santa Cruz, #SC-47701) and Protein G Dynabeads (Invitrogen, 10003D). DNA was purified using DNA purification columns (Zymo Research, D5205). Quantitative protein occupancy was performed by qPCR according to the manufacturer’s instructions (Jena Bioscience, PCR-369). Mean values for the analyzed regions were expressed as fold enrichment compared to a negative control without protein occupation (SOX2 gene). Statistically significant differences were analyzed using one-way ANOVA followed by the Fisher’s LSD post hoc test.

### ATAC-see

ATAC-see experiments were performed following the published protocol^31^. Briefly, following fixation and lysis, cells were incubated with the Tn5-ATTO-590N transposase (donated by Dr. Xingqi Chen) for 30 minutes at 37°C. Cells were washed at 55°C with PBS, SDS 0.01% and EDTA 50 mM to stop the tagmentation reaction, then counterstained with DAPI and mounted with Fluoromount. A negative control was conducted by adding EDTA 50 mM to the tagmentation mix to inhibit Tn5 activity. Image acquisition was performed in a LEICA TCS SP5 or Zeiss LSM800 microscope.

### Nascent RNA labeling

Nascent RNA imaging was performed using the Click-iT RNA imaging kit (ThermoFisher) following manufacturer’s instructions. Cells were fed with 5-ethynyl uridine 1 mM for 6 hours, fixated with formaldehyde 3.7%, and permeabilized with Triton X-100 0.5%. Click it reaction with Alexa Fluor 594 was carried out and nuclei were stained with Hoechst 33342. Image acquisition was performed in a Zeiss LSM800 microscope.

### Digital image processing

Image processing was performed using Fiji^72^. Nuclei were identified and segmented using a Gaussian filter on raw DAPI images and then applying an intensity threshold to generate nuclear outlines. All nuclear outlines were verified visually, and mitochondrial ATAC-see signal was filtered out using the nuclear outlines generated. A total of 50 nuclei were analyzed. Mean intensity per nucleus was measured and brightest regions were identified using an intensity threshold to retain only the pixels with intensity values higher than the mean.

Distance maps of individual nuclei were generated in Fiji to measure signal intensity as a function of radial distance to the nuclear periphery. Mean intensity was measured in a series of one-pixel-wide annuli from the outermost pixel to the nuclear center. Each channel was normalized to its mean intensity per nucleus and nuclear size was scaled in R. 50 individual nuclei were analyzed per experiment and average signal was calculated. Correlation analysis between signal intensity and distance from the nuclear rim was performed in R.

## Data accessibility

All data sets have been deposited in the GEO repository with accession number GSE206194. Access to the data sets will be provided upon paper acceptance in a peered review journal. For more information please contact the corresponding author.

## Acknowledgements

We thank Dr. Xingqi Chen for his generous donation of the Tn5-ATTO-590N enzyme. We thank Dr. Félix Recillas-Targa for kind donation of the HD3 erythroblast cell line. We thank the Molecular Biology Unit, the Microscopy Unit and the Bioinformatics Unit at the IFC and in particular to Dr. Ruth Rincón-Heredia and Dr. Augusto Cesar Poot-Hernández for advice on image processing and genomic data analysis respectively. We thank the Animal Facility at the IFC in particular Dr. Claudia Rivera-Cerecedo and Dr. Hector Malagón-Rivera. We thank the Supercomputing Unit at LANCIS and in particular to M.Sc. Rodrigo García-Herrera for access to the computer cluster. We thank Dr. Ricardo Saldaña-Meyer and Dr. Martín Escamilla Del Arenal for helpful discussion on the results obtained in this work.

## Authors Approval

All authors have seen and approved the manuscript. This manuscript hasn’t been accepted or published elsewhere.

---

**Extended data 1.**
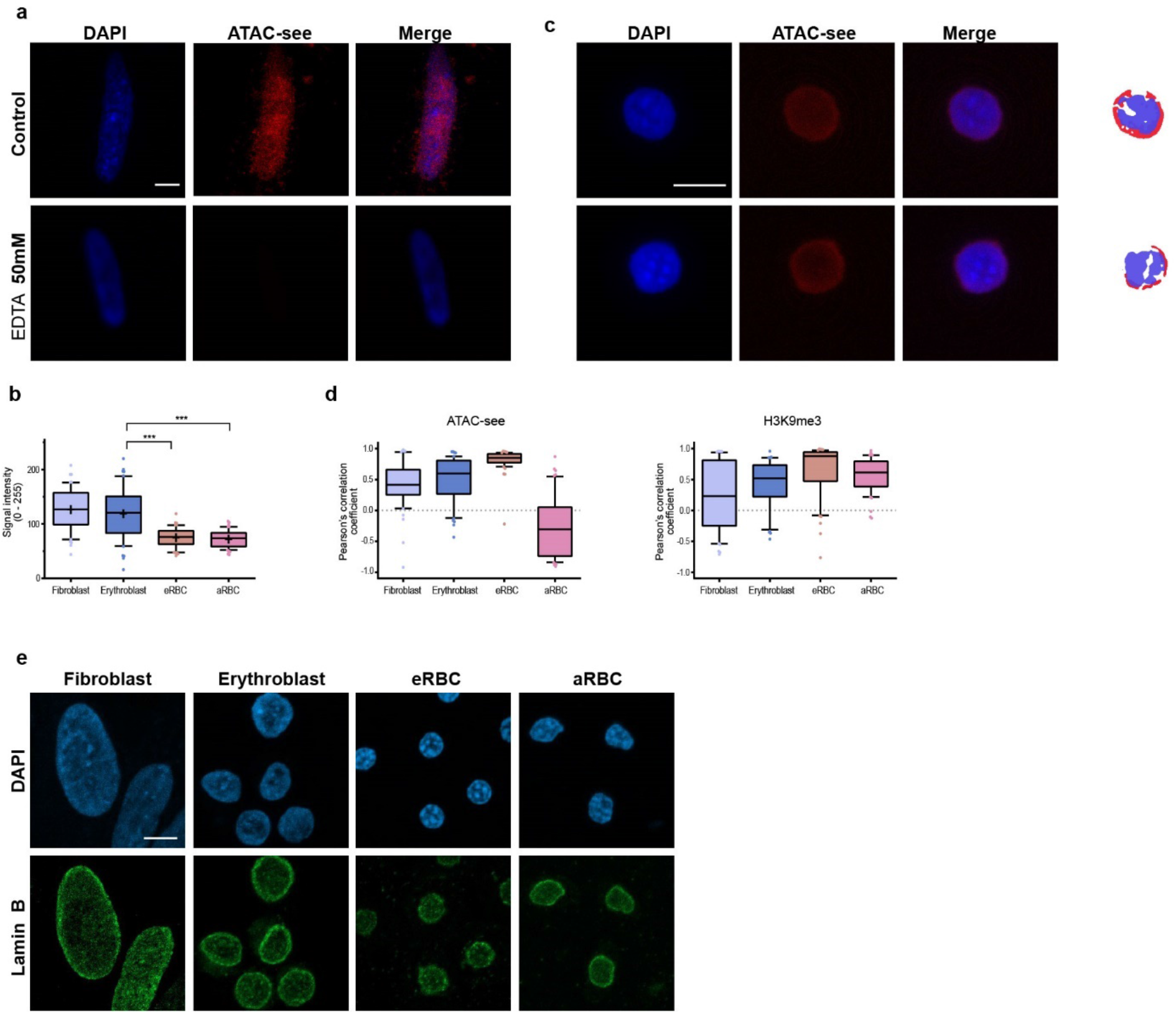
Validation of ATAC-see and chromatin inversion in aRBC. **a**, Imaging of open chromatin by ATAC-see (red) and DAPI (blue, nuclei) in chicken fibroblasts. A negative control of ATAC-see in presence of EDTA 50 mM shows no incorporation of the fluorophore. **b**, ATAC-see signal average intensity per nucleus in chicken erythrocytes. Boxplots and mean of measurements from 50 nuclei are shown. Asterisks indicate P values based on one-way ANOVA followed by Tukey’s post-hoc test. ***P < 0.001. **c**, Representative images of ATAC-see in aRBC. Brightest pixels enriched in ATAC-see (red) or DAPI (blue) are shown on the right. **d**, Pearson’s correlation score between ATAC-see or H3K9me3 signal intensity and the distance to the nuclear rim (n = 50) **e**, Fluorescence imaging of Lamin B1 (green) and DAPI. In every case, maximal projections of three-dimensional (3D) stacks are presented. Scale bar = 5 µm.

**Extended data 2.**
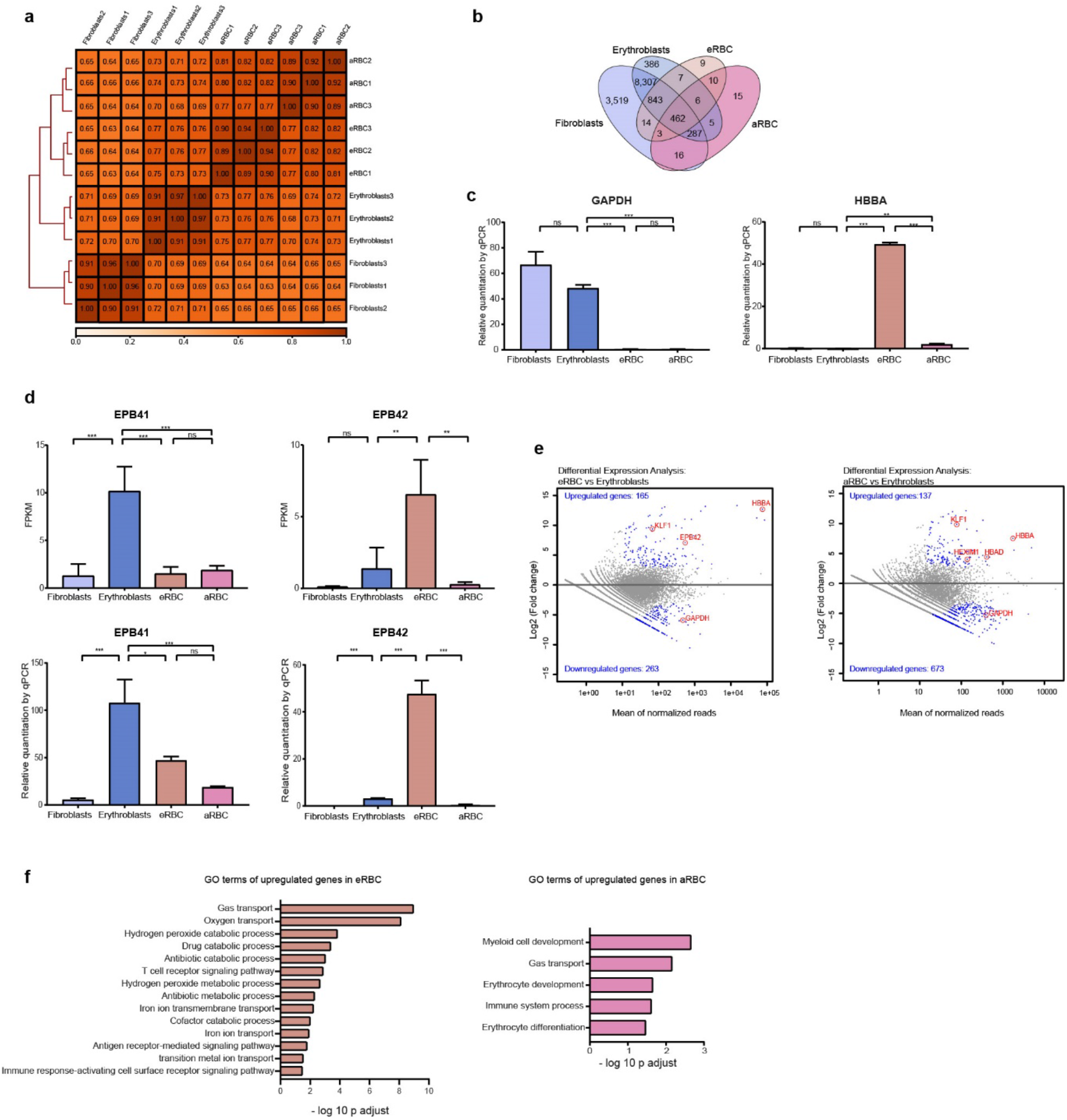
Characterization of the erythroid transcriptome. **a**, Spearman correlation heatmap of RNA-seq replicates. **b**, Venn diagram of expressed genes in fibroblasts and erythroid cells. **c**, Relative expression of GAPDH and HBBA measured by RT-qPCR. **d**, Expression levels of the erythroid genes EPB41 and EPB42 measured by RNA-seq (up) and RT-qPCR (down). Asterisks indicate P values based on one-way ANOVA followed by Tukey’s post-hoc test. **e**, MA plots of differentially expressed genes between eRBC and erythroblasts, and aRBC and erythroblasts. **f**, GO terms enriched in upregulated genes from **e**.

**Extended data 3.**
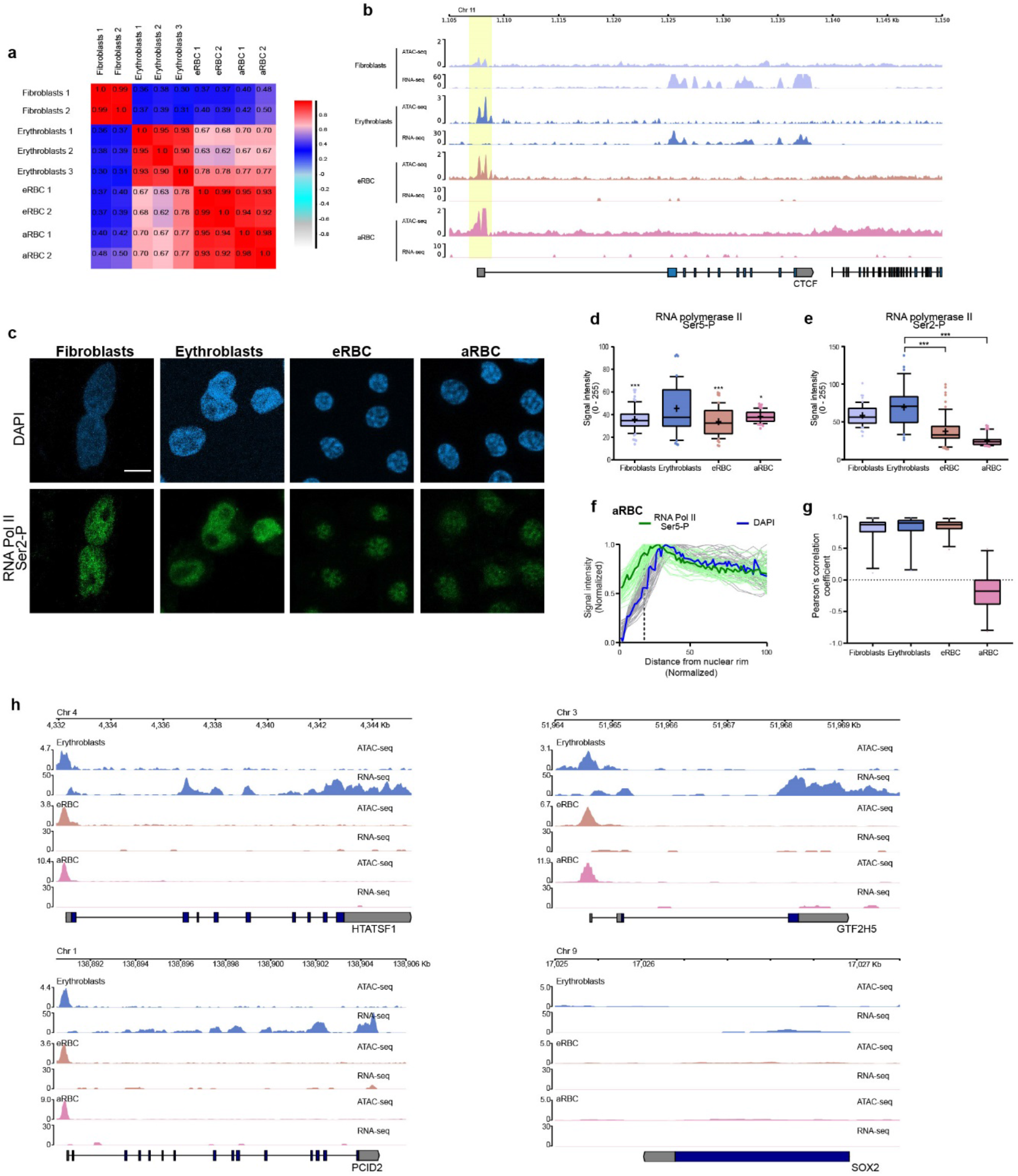
Chicken erythrocytes retain accessible paused promoters. **a**, Spearman correlation heatmap of ATAC-seq replicates. **b**, Example of the preserved accessibility of the promoter of CTCF despite no transcriptional activity detected by RNA-seq. **c**, Representative images of DAPI and RNA pol II Ser2-P immunofluorescence. Maximal projections of three-dimensional (3D) stacks are presented. Scale bar = 5 µm. Average signal intensity of RNA pol II Ser5-P (**d**,) and Ser2-P (**e**,) signal average intensity per nucleus in chicken erythrocytes. Boxplots and mean of measurements from 50 nuclei are shown. Asterisks indicate P values based on one-way ANOVA followed by Tukey’s post-hoc test. *P < 0.05; ***P < 0.001; ns not significant differences. **f**, RNA pol II Ser5-P and DAPI intensity is presented as a function of distance to the nuclear periphery. Individual tracks for 50 nuclei are presented in lighter colors and the mean intensity is presented in bold. **g**, Pearson’s correlation score between RNA pol II Ser5-P signal intensity and the distance to the nuclear rim (n = 50). **d**, ATAC-seq and RNA-seq tracks of HTATSF1, GTF2H5, PCID2, and SOX2.

**Extended data 4.**
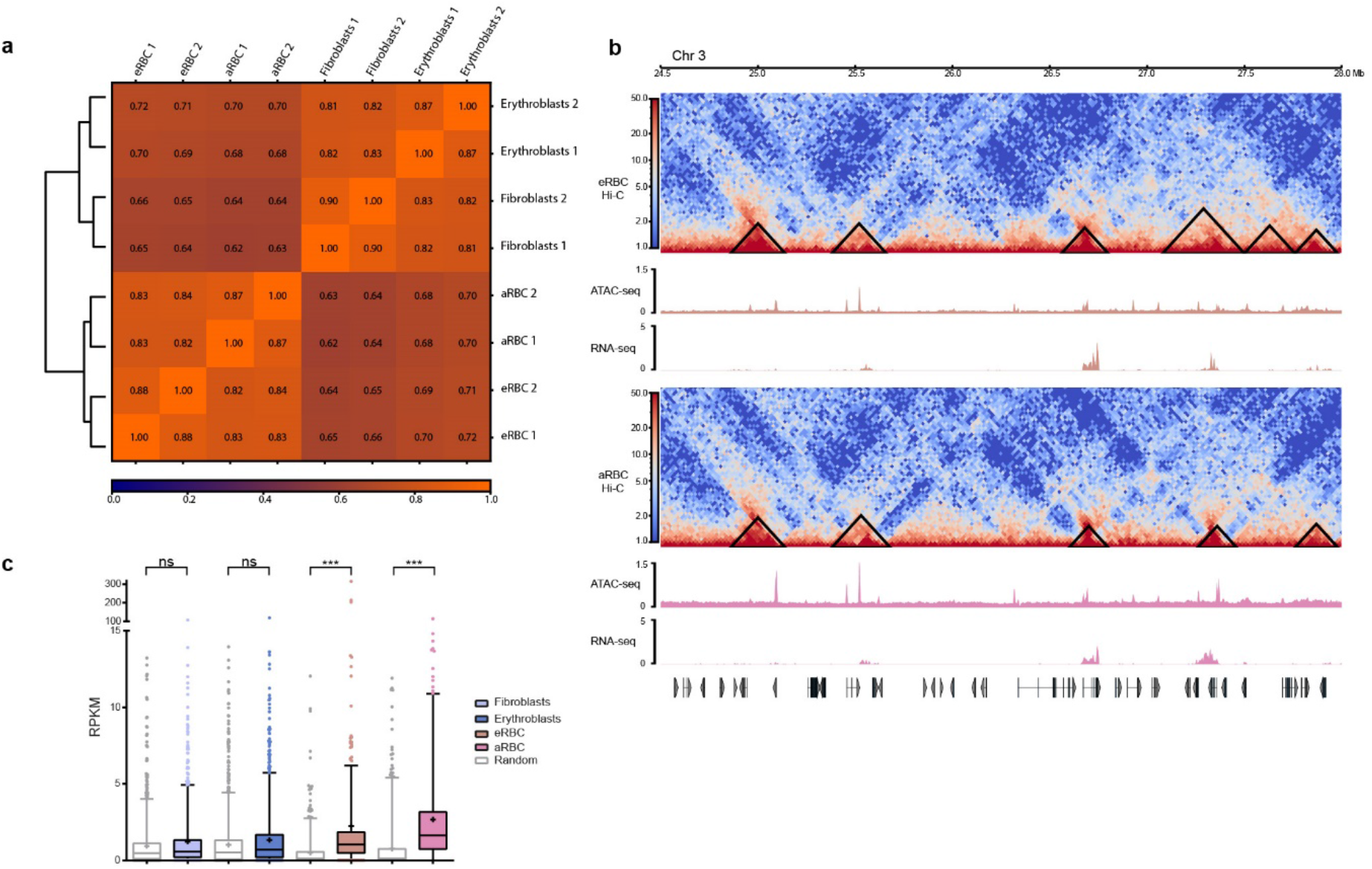
Erythroid mini domains retain local structure around open chromatin and paused genes. **a**, Pearson correlation heatmap between Hi-C replicates at 50 Kb resolution. **b**, Hi-C contact matrices from eRBC and aRBC (3: 24.5 – 28 Mb) and examples of mini domains identified. ATAC-seq and RNA-seq tracks are plotted below. **c**, RNA-seq RPKM count inside domains or a random control. Asterisks indicate P values based on one-way ANOVA followed by Kruskal-Wallis post hoc test. ns = P > 0.05; ***P < 0.001.

